# Spatiotemporally distinct responses to mechanical forces shape the developing seed of *Arabidopsis*

**DOI:** 10.1101/2023.08.21.554152

**Authors:** Amélie Bauer, Camille Bied, Adrien Delattre, Gwyneth Ingram, John F. Golz, Benoit Landrein

## Abstract

Organ morphogenesis depends on mechanical interactions between cells and tissues. These interactions generate forces that can be sensed by cells and affect key cellular processes. However, how mechanical forces contribute, together with biochemical signals, to the shaping of complex organs is still unclear. We address this question using the seed of *Arabidopsis as* a model system. We show that seeds first experience a phase of high anisotropic growth that is dependent on the response of cortical microtubule (CMT) to forces, which guide cellulose deposition according to shape-driven stresses in the outermost layer of the seed coat. However, at later stages of development, we show that seed growth is isotropic and depend on the properties of an inner layer of the seed coat that stiffens its walls in response to tension but has isotropic material properties. Finally, we show that the transition from anisotropic to isotropic growth is due to dampening of CMT responses to shape-driven stresses. Altogether, our work support that spatiotemporally distinct mechanical responses control the shape of developing seeds in *Arabidopsis*.

## Introduction

Like all biological organisms, plants are physical structures whose growth depends on the generation of forces (Landrein & Ingram, 2019). At the cellular scale, plant growth relies on the balance between the hydrostatic pressure of the cell (i.e., turgor) and the mechanical properties of the surrounding cell wall. Pressure promotes growth by generating tension in plant cell walls. When tension exceeds the yielding threshold of the walls, it induces their plastic deformation, and thus irreversible cell expansion (Boudaoud, 2010). Plant primary cell walls, which surround growing cells, are complex structures composed of stiff microfibrils of cellulose embedded in a gel-like matrix of pectins, of hemicelluloses, and of structural and regulatory proteins (Cosgrove, 2022). While the extensibility and the rigidity of the wall can be tuned by modifying the properties of any of these different components, its ability to stretch in a particular direction largely depends on the organization of the cellulose microfibrils, which are the main load-bearing structures of the wall. In this regard, cellulose microfibrils can be oriented in a particular direction to form crystalline arrays that resist stretching, forcing walls to expand perpendicularly to the main fibril axis (Cosgrove, 2022). Cellulose microfibrils are synthesized by multimeric transmembrane cellulose synthase complexes (CSCs) that move in the membrane following tracks composed of cortical microtubules (CMTs) (Paredez *et al*, 2006). By guiding the deposition of cellulose fibres, CMTs can therefore determine the mechanical anisotropy of the wall and control the main orientation of cell elongation.

At the organ scale, plant growth patterns depend on mechanical interactions between cells and tissues. This is due to the presence of the wall, which glues cells together so that differences in mechanical properties between adjacent cells or adjacent layers generate mechanical conflicts that can, when resolved, induce the emergence of complex shapes (Rebocho *et al*, 2017; Coen *et al*, 2004). In many aerial organs, it is believed that growth is promoted by the pressure of inner tissues but restricted by the mechanical properties of one or several outer layers, which is often the epidermis (Kutschera & Niklas, 2007). This model of growth, defined as the “epidermal growth control” theory, is supported by both genetics and mechanical studies (Kelly-Bellow *et al*, 2023; Savaldi-Goldstein *et al*, 2007; Verger *et al*, 2018; Asaoka *et al*, 2021; Vaseva *et al*, 2018). This model also implies that cells are submitted to specific patterns of mechanical forces that are dependent on their position within the organ but also on the overall shape of the organ (i.e. shape-driven stresses) (Hamant *et al*, 2008). Although the molecular mechanisms of mechanoperception are still largely unknown in the plant kingdom, accumulating evidence suggests that plant cells can sense mechanical signals, and use them as cues to assess shape changes occurring at the organ scale during morphogenesis (Landrein & Ingram, 2019). Mechanical forces have notably been shown to affect CMT organization, and thus cellulose deposition (Hamant *et al*, 2008; Sampathkumar *et al*, 2014; Robinson & Kuhlemeier, 2018; Hejnowicz *et al*, 2000), so that shape-driven stresses can affect anisotropic growth in a variety of plant organs (Trinh *et al*, 2021). However, to what extent mechanical signals contribute to the control of plant organ shape remains unclear, partly because CMTs can also respond to other signals, such as light and hormones (Sambade *et al*, 2012; Vineyard *et al*, 2013), but also because organ shape is the product of a complex set of mechanical interactions between cells and tissues, and does not only depend on the molecular mechanisms controlling cell growth anisotropy (Rebocho *et al*, 2017).

The seed of *Arabidopsis* is a powerful model system in which to study how mechanical interactions between layers contribute to plant organ growth. Seeds are composed of three genetically distinct compartments: the embryo, the endosperm and the surrounding seed coat (Ohto *et al*, 2007). As the growth of the seed precedes that of the embryo, it is thought to depend mainly on interactions between the endosperm and the seed coat. Accordingly, we recently provided evidence suggesting that seed growth is promoted by the pressure of the endosperm but restricted by the mechanical properties of an internal layer of the seed coat, called the adaxial epidermis of the outer integument (ad-oi layer) (Creff *et al*, 2023, 2015; Beauzamy *et al*, 2016). We demonstrated that this layer stiffens its internal cell wall (called wall 3) in response to the tension induced by endosperm expansion and that this stiffening is associated with the accumulation of demethylesterified pectins in the wall. Using modelling and experiments, we showed that mechanosensitive stiffening of seed coat walls could underlie seed size control in *Arabidopsis* and explain the counter-intuitive effect of pressure on growth (Creff *et al*, 2023). However, how this mechanism intersects with the mechanical feedback loop between CMTs and forces to determine seed shape, has not been investigated.

Here we show that seed growth patterns are not uniform and can be divided into two distinct phases: an early phase of rapid anisotropic growth followed by a phase of slower isotropic growth. We show that CMT responses to forces in the abaxial epidermis of the outer integument of the seed coat (ab-oi the outermost cell layer of the seed coat) drive the initial phase of anisotropic growth through the guided deposition of cellulose microfibrils in the outer wall of the seed. In contrast, we show that growth is restricted by the mechanical properties of wall 3 during the isotropic growth phase, that stiffens in response to forces, but should have isotropic material properties based on the organization of the CMTs in the adaxial epidermis of the outer integument (ad-oi). Finally, we show that the transition from anisotropic to isotropic growth does not depend on wall 3 stiffening but relies on a dampening on the response of CMTs to shape-driven stresses in the abaxial epidermis of the outer integument. Taken together, our work supports that the combined action and fine tuning of two mechanical responses occurring in two different layers of the seed coat and at different times, control both seed growth rate and seed growth anisotropy, thus determining both seed size and shape.

## RESULTS

### Seed growth is divided into an early phase of high anisotropic growth and a later phase of slow isotropic growth

To quantify the changes in size and shape that occur during seed development we sampled fruits daily following anthesis and extracted and optically cleared the developing seeds before imaging them by differential interference contrast (DIC) microscopy and measuring their size and shape on 2D images (Fig 1A-B). We observed, as we previously reported (Creff *et al*, 2023), that seed growth (measured as area) occurred within the first week following fertilization. During this time, seed size increased following a characteristic S-shaped pattern: growth was higher at the beginning of the growth phase but slowly decreased until it terminated at around 7 days post anthesis (Fig 1C). We then studied the shape changes that occurred during seed growth by measuring the seed aspect ratio (defined as the ratio between the length of the seed and its width (See Materials and Methods and Fig 1B)). To our surprise, we observed two distinct growth behaviours. During the first three days following anthesis, the aspect ratio, which in ovules, was close to 0.8, increased rapidly, to reach a value close to 1.7, due to the elongation of the seed along its future main axis. However, between 3 and 7 days post-anthesis, growth was much more isotropic as seeds expanded slightly more in width than in length (Fig 1D and Fig S1A-C). Taken together, these quantifications show that seed growth involves two distinct phases: an initial phase of rapid anisotropic growth followed by a phase of slow and almost completely isotropic growth.

**Fig. 1.**
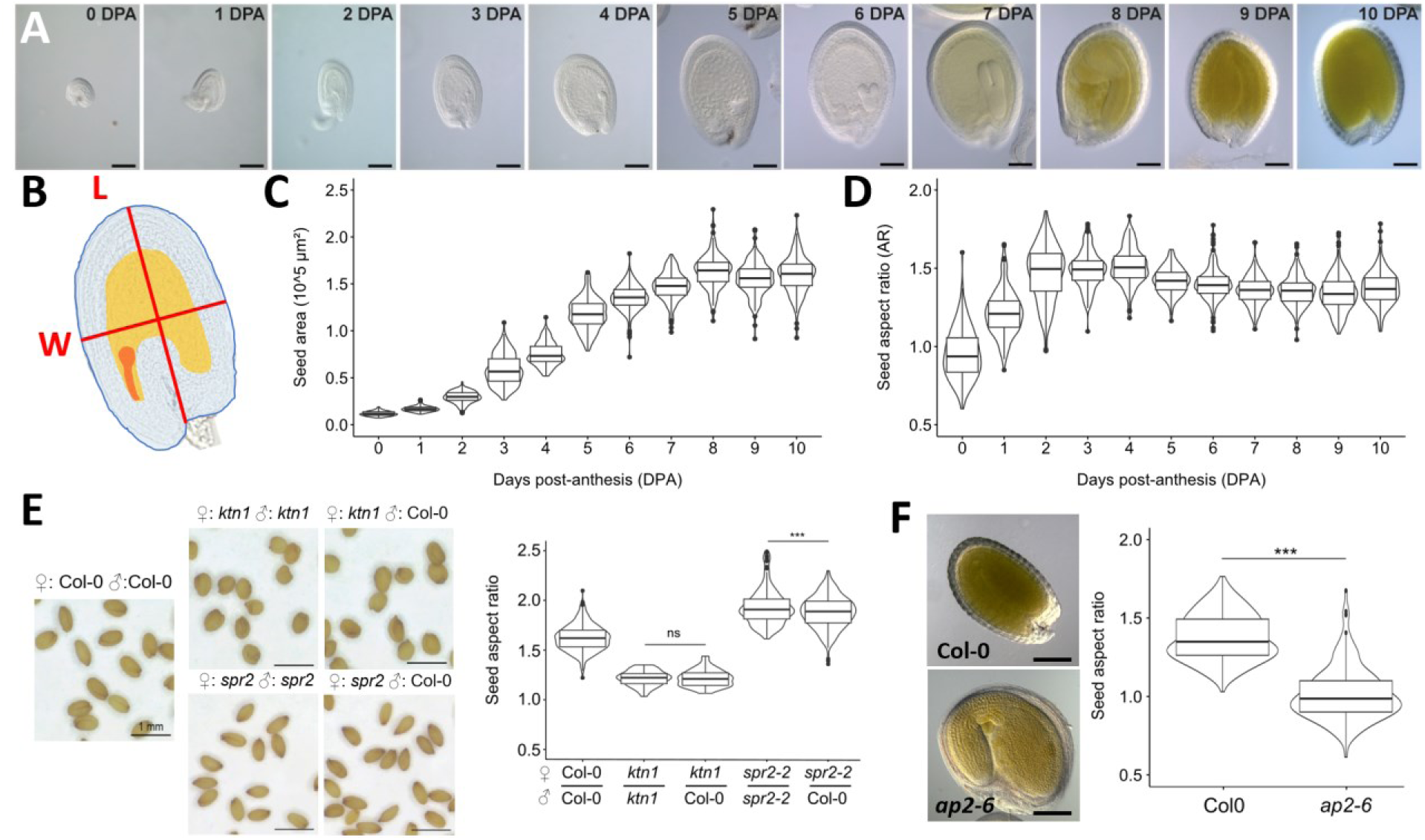
Seed growth anisotropy is not monotonous and is controlled by the seed coat outer integument. **A.** Representative WT seeds (Col-0 ecotype) from 0 to 10 days post-anthesis (DPA), scale bars: 100 µm. **B.** Seed aspect ratio is defined as the ratio between seed length (L, obtained by drawing a line from the pedicel of the seed to its tip that passes through the middle of the endosperm) and width (W, obtained by drawing a line perpendicular to the previous axis and passing through its centre). The seed coat is highlighted in blue, the endosperm in orange and the embryo in red. **C-D.** WT seed area (C) and aspect ratio (D) from 0 to 10 DPA, n=180-209 seeds per day, two independent experiments. **E.** Shape of mature seeds of *ktn1* and *spr2* mutants whose ovules were self-pollinated or pollinated with WT pollen, scale bars: 1 mm, n= 78-551 seeds, two independent experiments. Data were compared using bilateral Student tests. **F.** Shape of Col-0 and *ap2* seeds (pollinated with WT pollen) at 10 days post anthesis (DPA), n=162-230 seeds, two independent experiments. Data were compared using bilateral Student tests.

### Seed shape depends on the properties of the outer integument of the seed coat

We previously showed that seed size is the product of mechanical interactions between two seed compartments: the endosperm and the seed coat (Creff *et al*, 2023). In the proposed mechanical model, the endosperm is predicted to generate an isotropic force *via* its hydrostatic pressure whereas the cell walls of the seed coat could, if they have anisotropic mechanical properties, restrict growth in particular directions more than in others (Boudaoud, 2010; Creff *et al*, 2023). To test this hypothesis we examined the seed growth profile of mutant lines lacking functioning *KATANIN* (*KTN1*) and *SPIRAL* (*SPR2*), genes that encode microtubule associated proteins involved in CMT organization and responses to force. We observed that self-pollinated plants produced seeds with similar shape defects (rounder seeds for *ktn1* mutants and more elongated seeds for *spr2* mutants) as those arising from mutant ovules pollinated with WT pollen. Given that this latter class of seeds have phenotypically wildtype endosperm and embryos (being heterozygous), but a phenotypically mutant seed coat owing to its maternal origin (Fig 1E), our data suggest that seed shape mainly depend on the mechanical properties of the seed coat.

The seed coat is a maternal tissue that originates from differentiation of the ovule integuments and in *Arabidopsis*, is composed of five cell layers: the two outermost cell layers form the outer integument whereas the three innermost layers arise from the inner integument (Fig S1D) (Cucinotta *et al*, 2014). To test which layer of the seed coat determines seed shape, we characterized the shape of *apatela2* (*ap2*) mutant seeds, in which outer integument cells fail to differentiate (Ohto *et al*, 2005; Jofuku *et al*, 2005). We observed that *ap2* plants (crossed with WT pollen) were similarly round, showing that this gene controls seed growth anisotropy, and that seed shape depends on the differentiation of at least one layer of the outer integument (Fig 1F). These results are in agreement with the epidermal growth control theory that proposes organ growth to be restricted by the properties of outer cell layers (Kutschera & Niklas, 2007).

### Cells in the outer integument of the seed coat also show two distinct growth patterns

As our results so far suggest that seed shape depends on the mechanical properties of the outer integument, we compared the growth pattern of the cells within the outer integument during the anisotropic growth phase (from 1 to 2 DPA) and the isotropic growth phase (from 4 to 5 DPA). To do this, we isolated seeds expressing the ubiquitous membrane marker (*LTi6b-GFP*) from *in vitro-*grown fruits, by adapting a previously published protocol (Gooh *et al*, 2015), and imaged them by confocal microscopy at two different timepoints (0h and 24h). We then used the MorphoGraphX software and the level set method (LSM) to extract meshes of the outer-surface of both layers of the outer integument, segment the cells on these meshes and manually track them from one timepoint to the other (Barbier de Reuille *et al*, 2015; Kiss *et al*, 2017). The resulting growth maps of developping seeds enabled us to measure cell growth rate, growth anisotropy and division rate in the two layers of the outer integument (Fig 2A).

**Fig. 2.**
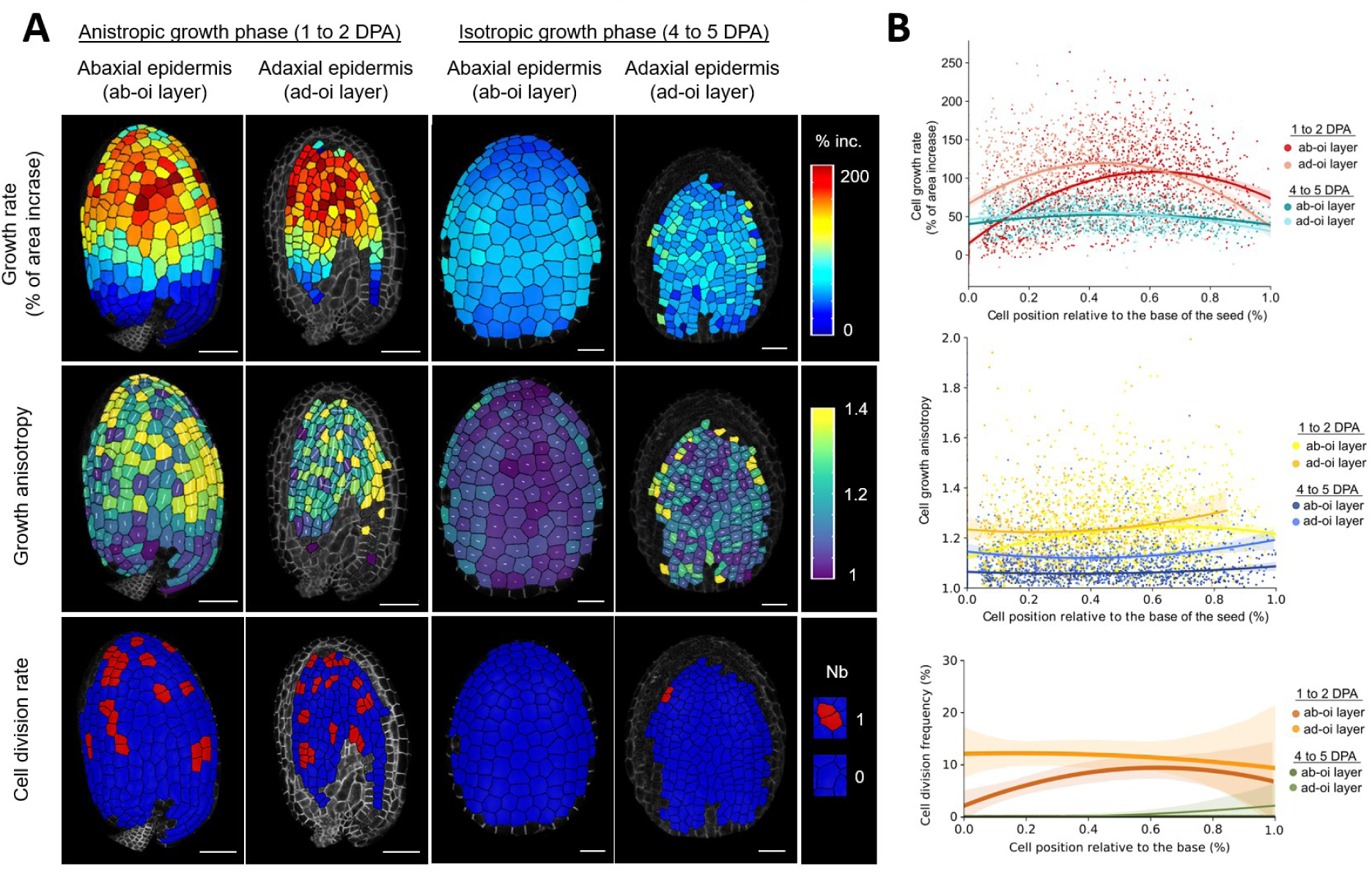
Quantifying cell growth in the two layers of the seed coat outer-integument. **A-B** Representative maps (A) and quantification (B) of cell growth rate, cell growth anisotropy and cell division in the abaxial (ab-oi) and adaxial (ad-oi) epidermis of the outer integument of the seed during the anisotropic growth phase (from 1 to 2 DPA (Days post-anthesis)) and during the isotropic growth phase (from 4 to 5 DPA). In the maps of growth anisotropy, the white lines overlaid on the heatmap represent the main axis of elongation of the cells. In the graphs in B, the coloured lines and dim bands show the mean and standard deviation as a function of the relative position with respect to the base of the seed. Scale bars: 20 µm, n=1015-1677 cells from 10-11 seeds, two independent experiments.

In agreement with our measurements at the organ scale (Fig 1C), we observed that cells in both layers of the outer integument grew faster during the anisotropic growth phase than during the isotropic growth phase (Fig 2). Furthermore, we also found that growth was not homogenous in the outer integument, especially during the anisotropic growth phase, where cells at the tip of the seed expanded much more than the cells at the base of the seed (Fig 2), which is in line with observations for other plant organs such as leaves and sepals (Hervieux *et al*, 2016; Kuchen *et al*, 2012). A characteristic feature of the anisotropic growth phase was the preferential elongation of outer integument cells parallel to the main axis of the seed, whereas during the isotropic growth phase there was no preferential direction of growth with cells acquiring a soap bubble organisation that is a defining mark of isotropic growth (Corson *et al*, 2009) (Fig 2 and Fig S2). Finally, we also observed that cells continued to divide during the anisotropic growth phase but not during isotropic growth phase so that they greatly enlarged over time, especially in the abaxial epidermis (Fig 2 and Fig S2). In summary, our quantitative analysis demonstrates that the early anisotropic growth of the seed is associated with the division and elongation of the cells of the outer integument that are located at the tip of the seed whereas the isotropic growth phase is associated with the slow and isotropic growth of all cells within the outer integument.

### The anisotropic growth phase depends on CMT organization in the abaxial epidermis of the outer integument

CMTs determine the main axis of cell elongation by controlling the oriented deposition of cellulose microfibrils in the wall (Paredez *et al*, 2006). In leaves, sepals and meristems, the CMTs facing the outermost cell wall of the epidermis, which is believed to be load-bearing, organize according to shape-driven stresses and are believed to control the anisotropic growth of these organs (Hamant *et al*, 2008; Hervieux *et al*, 2016; Sampathkumar *et al*, 2014; Zhao *et al*, 2020). However, in hypocotyls, it is the CMTs that face the inner side of the epidermis that are robustly oriented perpendicularly to the main axis of elongation (Crowell *et al*, 2011; Chan *et al*, 2011). This suggests that the inner wall of the epidermis is load-bearing and controls the elongation of this organ. This scenario is also consistent with computational simulations and experiments on hypocotyls (Robinson & Kuhlemeier, 2018; Verger *et al*, 2018) (Robinson 2018, Verger 2018).

To determine where seed growth anisotropy is controlled, we studied the organization of the CMTs in the two longitudinal faces of the two layers of the outer integument in the region that showed the highest growth rate in our maps (top half, Fig 2A and Fig S1D). We first compared the organization of the CMTs facing the outer side of the seed in the abaxial epidermis using an ubiquitous microtubule reporter (*p35S::MAP65-1-RFP*, (Creff *et al*, 2015)) to the organization of the CMTs facing both inner and outer side of the seed in the adaxial epidermis using a reporter we developped for this purpose (*pELA1::MAP61-1-mCitrine*, see Materials and Methods). We used the Fibritool plugin in ImageJ to determine the main orientation and the degree of organization of the CMTs near each cell face (Boudaoud *et al*, 2014). We observed that the CMTs facing the outer-face of the seed in the abaxial epidermis were highly organized and preferentially oriented perpendicularly to the main axis of the seed (Fig 3A-C). In contrast, we observed that the CMTs facing either side of the seed in the adaxial epidermis were less organized and were not aligned in any particular direction (Fig 3A-C). These data suggest that the CMTs of the abaxial epidermis of the outer integument determine the axis of seed elongation during the anisotropic phase, likely by driving cellulose deposition perpendicular to the main axis of the seed.

**Fig. 3.**
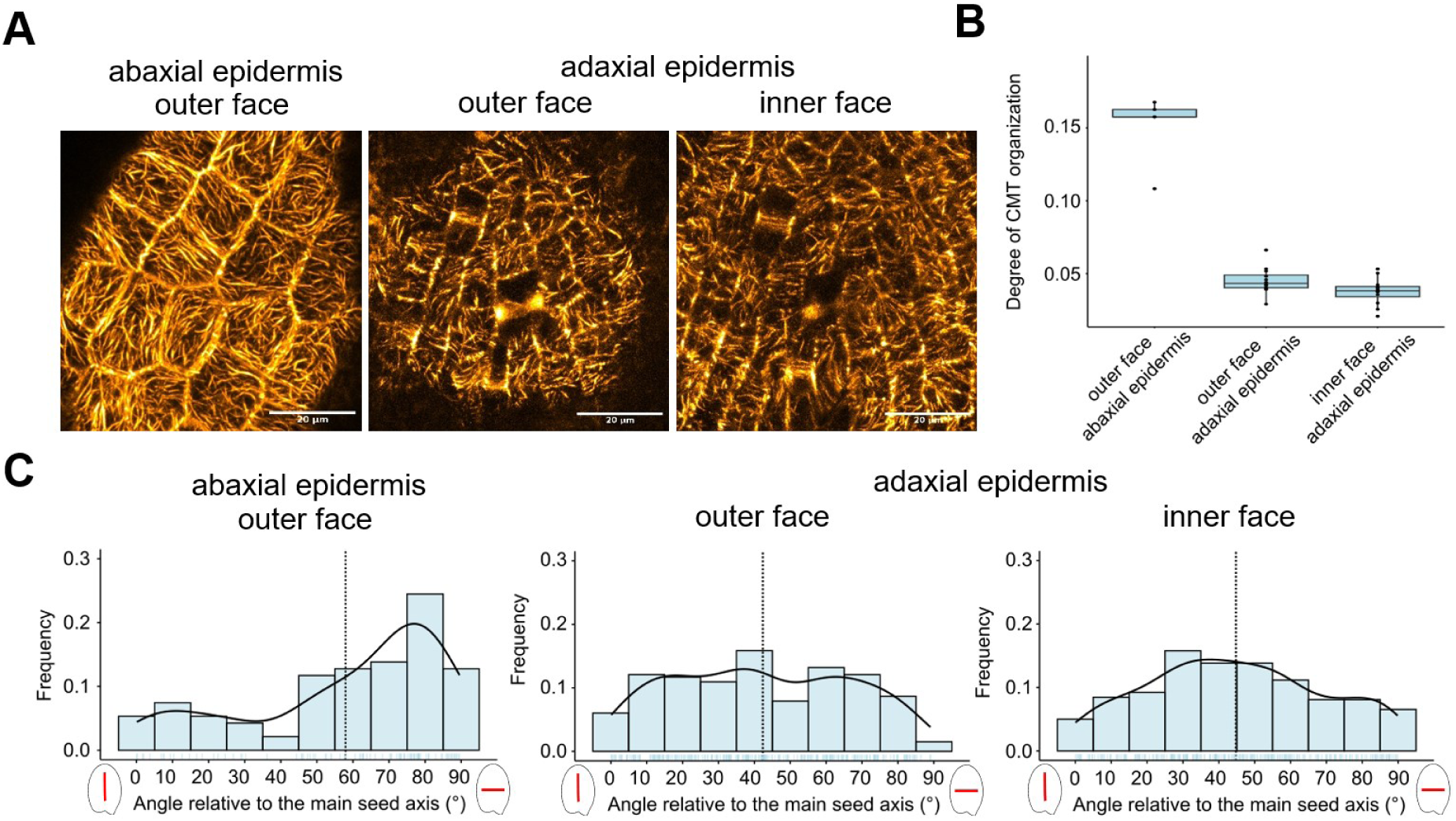
The CMTs of the abaxial outer integument epidermis control the initial phase anisotropic growth. **A.** Visualization of the organization of the CMTs in the outer face of the abaxial outer integument epidermis (imaged using a *p35S::MAP65-1-RFP* reporter) and in both faces of the adaxial outer integument epidermis (imaged using a *pELA1::MAP65-mCitrine* reporter) during the anisotropic growth phase (2 days post-anthesis). Scale bars: 20 µm. **B and C**. Degree of organization (B) and mean orientation relative to the main axis of the seed (C) of the CMTs in the outer face of the abaxial outer integument epidermis and in the two faces of the adaxial outer integument epidermis during the anisotropic growth phase (2 days post-anthesis), n= 94 to 265 cells from 5 to 12 seeds, two independent experiments. In the frequency plots in C, the bold lines show the trending curves and the dotted lines show the mean orientations of the CMTs relative to the main seed axis.

We also analysed the organization of the CMTs facing the outer-side of the seed in the abaxial epidermis with another ubiquitous reporter (*p35S::TUA6-GFP*), and facing both side of the seed using a reporter that is specifically expressed in the abaxial epidermis (*pPDF1::mCitrine-MBD* (Malivert *et al*, 2021)). As with the *p35S::MAP65-1-RFP* reporter, we observed that the CMTs located to the outer face of the abaxial epidermis preferentially aligned perpendicularly to the main axis of the seed (Fig S3A). With the *pPDF1::mCitrine-MBD* reporter, we also observed that the CMTs of the abaxial epidermis facing the inner side of the seed were preferentially oriented, and organized similarly, to the ones facing the outer side of the seed (Fig S3D). Collectively, these observations provide compelling evidence for the CMTs of the abaxial outer integument layer (in the outer face, the inner face or in both faces) controlling the elongation of the seed during the anistropic growth phase.

When comparing the reporters, we also observed that the CMTs were slightly less organized in the *p35S::TUA6-GFP* reporter and dramatically less organized in the *pPDF1::mCitrine-MBD* reporter than they were in the *p35S::MAP65-1-RFP* reporter (Fig S3B), suggesting that the presence of the reporter may affect the dynamics of the CMTs in the abaxial epidermis (and notably their bundling, Fig S3A). To test whether this observed effect of the reporters on CMT organisation affects the anisotropic growth of the seed, we compared the shape of mature seeds expressing these three reporters to seeds derived from control plants expressing a nuclear reporter in the abaxial epidermis of the outer integument (*pPDF1::CFP-N7* (Landrein *et al*, 2015)). This analysis revealed that the seeds expressing the *p35S::MAP65-1-RFP* reporter were 12% more elongated than control seeds while the seeds of the *pPDF1::mCitrine-MBD* reporter were 10% less elongated than control seeds (Fig S3C), consistent with the differences in CMT organization between these lines (Fig S3B). Given the influence of the CMT reporters on CMT organization and seed shape, our studies suggest that quantifications of CMT organization should be interpreted with caution, as they may depend on the nature of the CMT reporter, while the measurements of CMT orientation seem to be reporter independent.

### CMT response to forces in the abaxial epidermis controls the elongation of the seed during the anisotropic growth phase

Mechanical constraints affect CMT organization in a variety of plant organs (Landrein & Ingram, 2019), including seeds (Creff *et al*, 2015). We confirmed these findings by showing an increased organization of the CMTs facing the outer side of the seed in the abaxial epidermis of *p35S::MAP65-1-RFP* expressing seeds following a 24h compression of the fruit (Fig 4A). To study whether tissue shape is sufficient to prescribe a pattern of stress consistent with CMT alignment, we generated curvature maps of the surface of the seed so that we could correlate the orientation of the CMTs in the outer-face of the abaxial epidermis (using the ubiquitous microtubule reporter *p35S::MAP65-1-RFP*) with the local curvature of the seed (Fig 4B). We observed that CMTs preferentially align along the main axis of curvature of the seed in the outer-face of the outer integument, especially at the flanks and at the tip of the seed, which are its most curved areas and should thus be submitted to higher streses (Fig 4B).

**Fig. 4.**
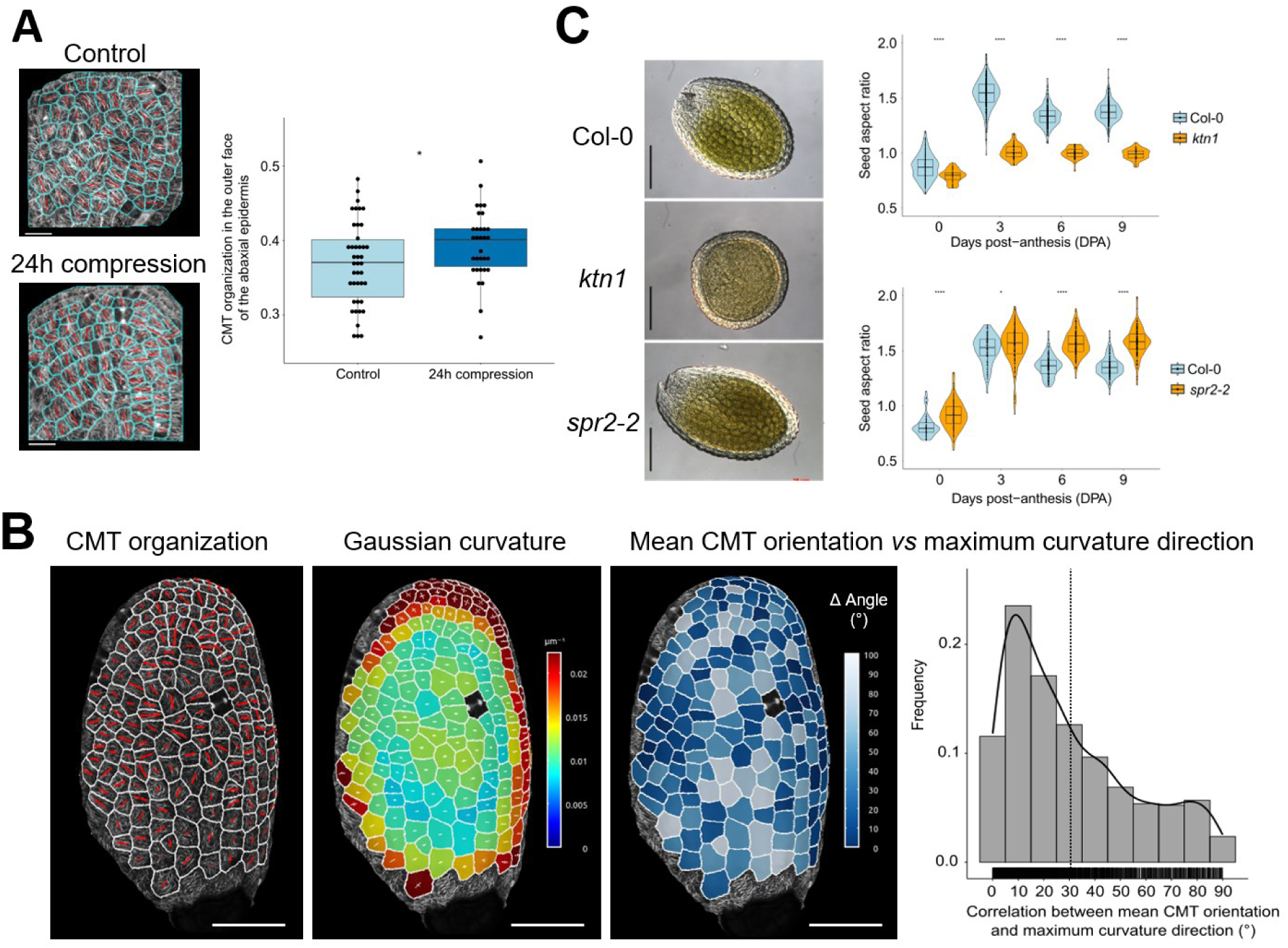
CMT response to forces in the abaxial outer integument epidermis drives seed growth anisotropy. **A.** Organization of the CMTs (*p35S::MAP65-1-RFP* reporter) in the outer face of abaxial outer integument epidermis during the anisotropic growth phase (2 days post anthesis (DPA)) in control seeds or in seeds whose fruits were compressed for 24h with a microvice (following the protocol of Creff *et al*, 2015), n=1512 to 2319 cells from 23 to 32 seeds, five independent experiments. In the pictures on the left, the orientation of the red bars shows the mean orientation of the CMTs in each cell and its length shows its degree of organization. Data were compared using a bilateral Student test. **B.** Correlation between the mean orientation of the CMTs in the outer face of abaxial epidermis (imaged using the *p35S::MAP65-1-RFP* reporter) and the Gaussian curvature of the seed during the anisotropic growth phase (2 DPA), n= 2252 cells from 19 seeds, five independent experiments. In the CMT picture, the orientation of the red bars shows the mean orientation of the CMTs in each cell and its length, their degree of organization. In the heatmap of Gaussian curvature, the orientations of the two perpendicular white bars represent the axes of maximum and minimum curvature in each cell and their lengths the degree of curvature in each of these two directions. In the frequency plot, the bold line shows the trending curve and the dotted line shows the mean angle between CMT orientation and maximum curvature direction. **C.** Seed shape defects of the mutants of CMT organization and response to forces: *katanin* (*ktn1*) and *spiral 2* (*spr2*). Scale bars: 100 µm, n= 28 to 152 seeds per day, two independent experiments. Data were compared using a bilateral Student tests.

To confirm that CMT responses to shape-driven stresses promote seed elongation through the guided deposition of cellulose microfibrils in the wall, we analysed the seed-growth defects of two mutants for genes involved in the control of microtubule organization and responses to forces: *katanin* (*ktn1*) and *spiral2 (spr2*), two mutants for genes involved in anchoring CSCs to CMTs: *cellulose synthase interacting protein 1* (*csi1*) and *tetratricopeptide thioredoxin-like 1 and 3 (ttl1 ttl3*), and one mutant for a gene that is part of the primary CSC: *procuste1/cellulose synthase 6* (*prc1-1*) (Hervieux *et al*, 2016; Uyttewaal *et al*, 2012; Kesten *et al*, 2022; Fagard *et al*, 2000; Bringmann *et al*, 2012). We observed seed shape defects that correlated with the function of the associated genes for all of these mutants (Fig 4C and Fig S4): reducing CMT responses to force in *ktn1*, CMT guidance of the CSCs in *csi1* and *ttl1 ttl3*, or cellulose synthesis in *prc1-1* led to reduced growth anisotropy and the production of rounder seeds (Fig 4C and Fig S4); in contrast, increasing CMT organization and response to forces in *spr2* led to increased growth anisotropy and the production of more elongated seeds (Fig 4C). In summary, the analysis of the seed shape defects in these mutants supports that CMT response to forces drives the anisotropic growth of the seed through the guided deposition of cellulose microfibrils in the wall.

### The adaxial epidermis of the outer integument controls isotropic growth at late stages of seed development

Having shown that CMT responses to forces in the abaxial epidermis of the outer integument promotes growth anisotropy at early stages of seed development, we then asked what controls the isotropic growth of the seeds at later stages of development (from 3 to 7 DPA). We recently showed that endosperm pressure directly promotes growth but indirectly inhibits it through the tension it generates in the seed coat, which promotes stiffening of the walls of the inner face (wall 3) of the adaxial outer integument epidermis (Creff *et al*, 2023). Given that growth becomes completely isotropic around the time at which wall 3 stiffens (at 3DPA, Fig 1D), we considered whether the observed isotropic growth of the seed might arise from the transfer of load-bearing stresses from the outer walls of the abaxial epidermis (wall 1 or 2) with their anisotropic material properties, to the walls of the inner face of the adaxial epidermis (wall 3) that have isotropic material properties.

To test this, we observed the organization of the CMTs in the inner and outer faces of the adaxial epidermis of seeds expressing the *pELA1::MAP65-1-mCitrine* reporter during the isotropic growth phase. Contrary to our observations made during the anisotropic growth phase (Fig 3), CMTs facing the outer side of the seed in the adaxial epidermis appeared organized and preferentially oriented perpendicularly to the axis of the seed, whereas those facing the inner side of the seed (i.e., facing wall 3), were not particularly organized and appeared to be slightly preferentially oriented in parallel to the main axis of the seed (Fig 5A-C). These observations are consistent with the hypothesis that wall 3, which is expected to be the main load-bearing wall during the isotropic growth phase (Creff *et al*, 2015, Creff, Ali *et al*, 2023), has isotropic material properties and controls isotropic growth at later stages of development.

**Fig. 5.**
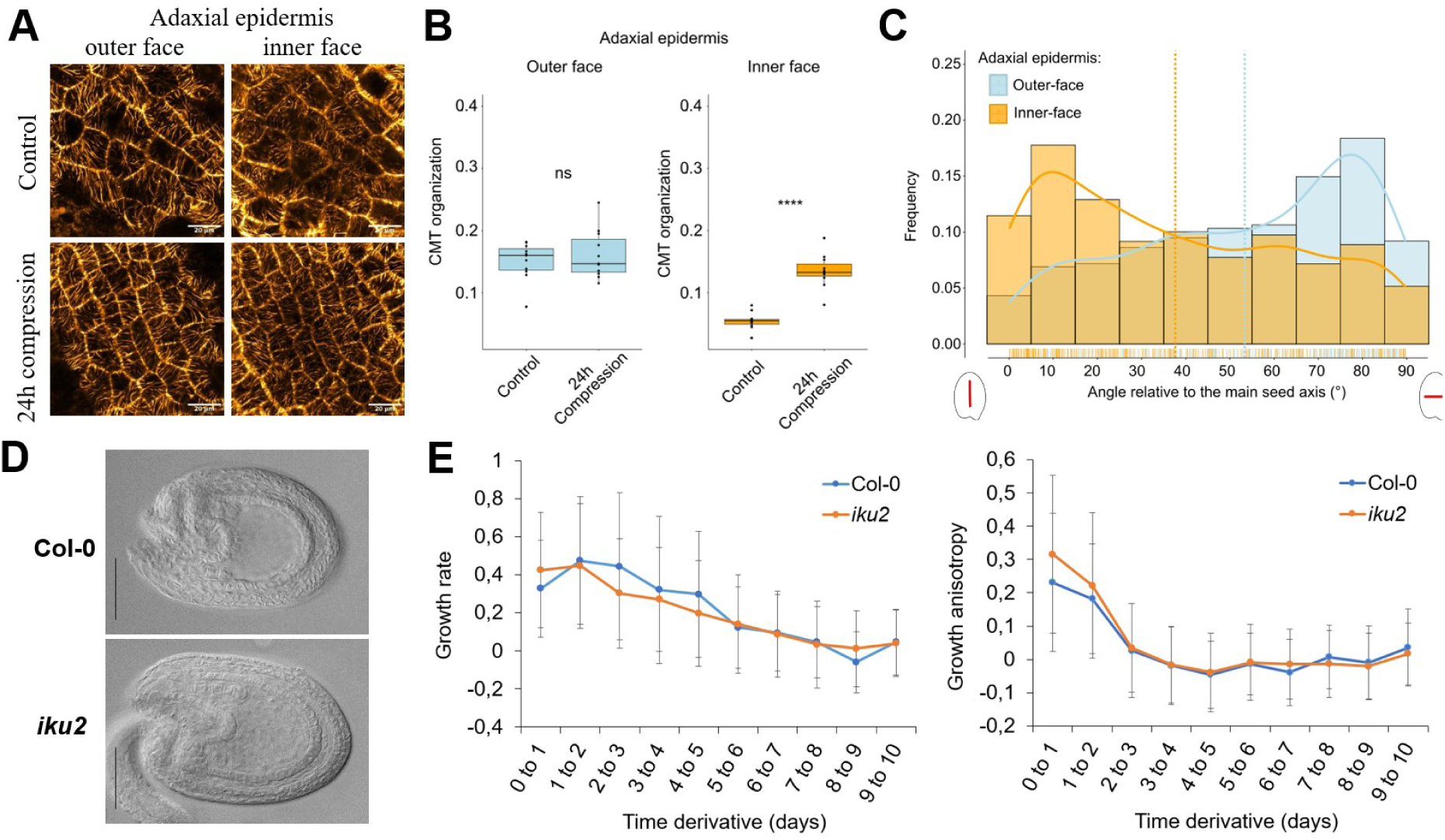
Wall 3 stiffening controls isotropic seed growth at late stages of development but is not responsible for the transition from anisotropic to isotropic growth. **A-B** Representative images (A) and quantification of the degree of organization (B) of the CMTs (imaged using the *pELA1::MAP65-1-mCitrine* reporter) facing the inner and the outer face of the seed in the adaxial outer integument epidermis during the isotropic growth phase (5 days post-anthesis (DPA)) in control seeds or in seeds whose fruits were compressed for 24h. Scale bars: 20 µm, n= 183 to 299 cells from 11 seeds per condition, two independent experiments. Data were compared using bilateral Student tests. **C.** Mean orientation relative to the main axis of the seed (C) of the CMTs facing the inner or the outer side of the seed in the adaxial outer integument epidermis during the isotropic growth phase (5DPA), n= 395 cells from 19 seeds, three independent experiments. The bold lines show the trending curves and the dotted lines show mean orientations relative to the seed axis. **D.** Representative Col-0 and *iku2* mutant seeds at 2 DPA, scale bar: 100 µm. **E.** Mean growth rate and anisotropy of developing WT and *iku2* mutant seeds from 0 to 10 days post-anthesis (DPA) obtained from the measurements of seed size and aspect ratio presented in Fig S5, n=207-313 seeds per day per genotype, three independent experiments.

The isotropic CMT organization we observed in the inner face of the adaxial epidermis could be associated with reduced mechanosensitivity of the CMT, or it could be the consequence of a reduction in stress levels or/and in stress anisotropy in this layer. To discriminate between these two hypothesis, we studied the response of the CMTs to the application of mechanical forces in both sides of the adaxial epidermis. We observed that a 24h compression of the fruit led to a strong increase of the organization of the CMTs facing the inner-side of the seed, which were initially not particularly organized, but not of the CMTs facing the outer-side, which were already organized (Fig. 5A-B). This demonstrates that the CMTs of the inner face of the adaxial outer integument epidermis still have the ability to respond to changes in stress pattern, and rather support that these CMTs are not organized either because stress is isotropic in this layer or because stress levels are too low to induce their organization.

To test if the transition from anisotropic to isotropic growth is a consequence of the stiffening of wall 3, we looked more closely at the correlation between seed growth rate and growth anisotropy, which were obtained by measuring of seed size and aspect ratio. We reasoned that if these two parameters are correlated, then they might both depend on wall 3 stiffening, not only in the wild type but also in *haiku2* (*iku2*), an endosperm-defective mutant in which higher endosperm pressure leads to more tension in the seed coat, inducing a precocious stiffening of wall 3 and an early restriction of growth (Creff *et al*, 2023; Garcia *et al*, 2003). In the wild type, we observed that both seed growth rate and seed growth anisotropy decreased over time but that seed growth anisotropy decreased earlier (peaking from 0 to 1 DPA) than seed growth rate (peaking between 2 to 3 DPA) (Fig 5E and Fig S5). In the *iku2* mutant, seeds elongated more rapidly than WT seeds between 0 to 1 DPA. This is expected as the seed coat of this mutant is under more tension than in WT seeds, which should enhance CMT organization during the anisotropic growth phase (Fig 5D-E and Fig S5). However, after 1 DPA, we observed that growth anisotropy was strongly decreased in *iku2*, reaching similar levels to those observed of WT, and that this reduction growth anisotropy also precedes the earlier restriction of growth observed in this mutant. These observations do not invalidate the idea that wall 3 is load-bearing and controls isotropic growth at later stages of development, but they suggest that the reduction of seed growth anisotropy observed between 1 and 3 DPA is a direct consequence of wall 3 stiffening.

### The transition from anisotropic to isotropic growth is associated with a disorganization of the CMTs in the outer face of the abaxial outer integument epidermis

If it is not a direct consequence of wall 3 stiffening, we reasoned that the transition from anisotropic to isotropic growth could be associated with a dampening of the CMT response to forces in the abaxial epidermis. To test this possibility, we compared the organization of the CMTs facing the outer face of the seed during both anisotropic and isotropic growth phase. Observations of seeds expressing of *p35S::MAP65-1-RFP* and *p35S::TUA6-GFP* reporters, but not of seeds expressing the *pPDF1::mCitrine-MBD* reporter (whose CMTs are not very organized at 2DPA, Fig S3B), supported that the CMTs facing the outer face of the abaxial epidermis become disorganized during the isotropic growth phase when compared to the anisotropic growth phase (Fig 6A and Fig S6A). Furthermore, analysis all three reporters revealed that CMTs were slightly less preferentially oriented perpendicularly to the main axis of the seed and in seeds of the *p35S::MAP65-1-RFP* reporter, that they were also slightly less prone to orient according to shape-driven stresses (Fig 6A-B and Fig S6A). These results indicate that the transition from anisotropic to isotropic growth is correlated with a reduced organization of the CMTs, although they can still, to some extent, orient according to shape-driven stresses.

**Fig. 6.**
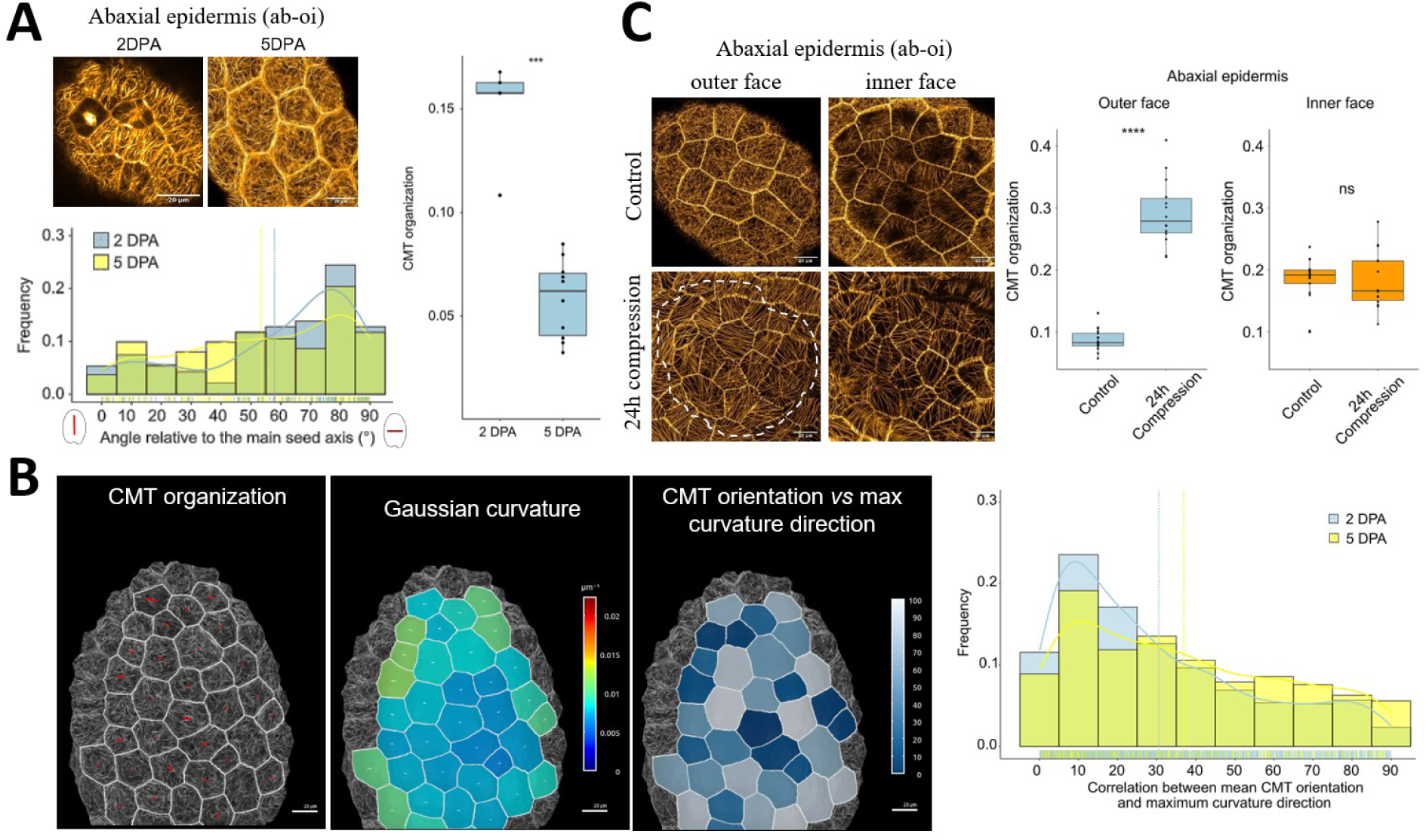
The CMTs facing the outer face of the abaxial outer integument epidermis are still mechanosensitive but become disorganized during the transition to isotropic growth. **A.** Comparison between the organization and the orientation relative to the main axis of the seed of the CMTs (images using the *p35S::MAP65-1-RFP* reporter) facing the outer side of the seed in the abaxial outer integument epidermis during the anisotropic growth phase (2 Days post-anthesis, DPA)) and the isotropic growth phase (5 DPA). Scale bars: 20 µm, n= 94 to 162 cells from 5 to 10 seeds, two independent experiments. In the frequency plot, the bold lines show the trending curves and the dotted lines show mean orientations relative to the seed axis. **B.** Comparison between the orientation of the CMTs (imaged using the *MAP65-1-RFP* reporter) facing the outer side of the seed in the abaxial epidermis and the local curvature of the seed during the anisotropic growth phase (2 DPA, see Fig. 4B for representative pictures) and the isotropic growth phase (5 DPA), scale bars: 20 µm, 2 DPA: n= 2252 cells from 19 seeds, five independent experiments, 5 DPA: n= 303 cells from 8 seeds, two independent experiments. In the CMT picture, the orientations of the red bars show the mean orientation of the CMTs in each cell and their lengths, their degree of organization. In the heatmap of Gaussian curvature, the orientations of the two perpendicular white bars represent the axes of maximum and minimum curvature in each cell and their lengths the degree of curvature in each of these two directions. In the frequency plot, the bold lines show the trending curves and the dotted lines show the mean angles between CMT orientation and maximum curvature direction. **C.** Effect of a 24h compression of the fruit on the mean degree of organization of the CMTs (imaged using the *pPDF1::Citrine-MBD* reporter) facing the inner or the outer side of the seed in the abaxial outer integument epidermis, scale bars: 20 µm, n= 273 to 293 cells from 14 to 18 seeds, two independent experiments. Data were compared using bilateral Student tests.

We also compared the organization and response to forces of CMTs in the two faces of the abaxial epidermis using the *pPDF1::mCitrine-MBD* reporter to test if the reduced CMT organization in the outer face of the abaxial outer integument epidermis could be the consequence of reduced mechanosensitivity. Once again, we observed that the CMTs facing the outer side of the seed, which were normally disorganized, strongly responded to seed compression, while the CMTs facing the inner side of the seed, which were already organized, did not show any increased organization in response to seed compression (Fig 6C). We also performed ablation studies to determine whether CMTs in the outer face of the abaxial epidermis respond to other types of mechanical perturbations (Hamant *et al*, 2008). After 6 hours, we observed a circumferential reorientation of the CMTs around the ablation site (Fig S6B). These data support that the transition from anisotropic to isotropic growth is not associated with a reduced mechanosensitivity of the CMTs in the outer face of the abaxial outer integument epidermis but could be due to a reduction in stress levels or/and anisotropy.

## Discussion

Seed growth depends on mechanical interactions between the zygotic endosperm and the surrounding maternally-derived seed coat. We previously showed that endosperm pressure both promotes and restricts seed growth through the tension it generates in the seed coat, which induces wall stiffening in the adaxial epidermis of the outer integument (Creff *et al*, 2023). Here, our results extend this model by showing that endosperm pressure controls seed shape by triggering sequential and different responses in neighbouring cell layers of the outer integument. At early stages of development, CMTs respond to shape-driven stresses in the abaxial epidermis, resulting in the elongation of the seed along its main axis. At later stages of development, endosperm pressure induces stiffening of the wall facing the inner side of the adaxial epidermis (wall 3). This wall subsequently becomes load bearing, and based on the organization of its CMTs, controls growth isotropically. Finally, our results show that the transition from anisotropic to isotropic growth does not depend directly on the mechanosensitive stiffening of wall 3, but involves a dampening of the response of the CMTs facing the outer side of the seed in the abaxial epidermis to shape-driven stresses, even-though these CMTs remain mechanosensitive.

Although our results show that CMT responses to forces in the abaxial epidermis of the outer integument promote the early anisotropic growth of the seed, previous work has suggested that the feedback between CMTs and forces amplifies growth anisotropy but is not necessarily its initial trigger. For instance, the elongation of developing hypocotyls is promoted by CMT response to forces and subsequent cellulose deposition according to tissue-stresses, but it is initiated as a result of an asymmetric distribution of methylesterified and demethylesterified pectins between transversal and longitudinal walls in the epidermis (Peaucelle *et al*, 2015). In pavement cells, differences in mechanical properties between contiguous walls, also associated with local pectin accumulation, trigger the initiation of the lobes through buckling, which are then amplified through CMT responses to forces (Majda *et al*, 2017; Sampathkumar *et al*, 2014). In lateral roots, elongation depends on CMT guidance of cellulose deposition but also on an independent membrane trafficking pathway which occurs at cell edges and requires the activity of the small GTPase Rab-A5c (Kirchhelle *et al*, 2019). Finally, in leaves and sepals, CMT responses to force are necessary for the maintenance of organ flatness, but flatness also requires an initial degree of organ asymmetry generated by the influence of margin-specific genes on early primordium growth (Zhao *et al*, 2020). The ovules of *Arabidopsis* have the shape of a flatten ovoid (Vijayan *et al*, 2021), which intuitively, should elongate in a direction opposite to that which is actually observed in fertilized seeds. Our results show that early developing seeds exhibit an apicobasal growth gradient. Whether this initial asymmetry in growth, acting on the ovoid shape of the ovule, is sufficient to generate a pattern of mechanical forces that induces the elongation of the seed along its main axis, or requires the activity of specific growth regulators such as *APETALA2*, remains to be demonstrated and will necessitate the generation of mechanical models of developing seeds that are based on realistic templates.

The ability of CMTs to respond to forces and to align along the main axis of tensile stress has been observed in a variety of plant organs, such as leaves, hypocotyls, sepals and meristems, and even in confined protoplasts (Hamant *et al*, 2008; Zhao *et al*, 2020; Sampathkumar *et al*, 2014; Colin *et al*, 2020; Robinson & Kuhlemeier, 2018). It has thus been hypothesized that CMTs could, by themselves, act as tension sensors (Hamant *et al*, 2019). However, this view might be oversimplistic as we know that CMTs can also respond to other signals, such as light and hormones (Sambade *et al*, 2012; Vineyard *et al*, 2013), and that their organization therefore does not always correlate with predicted stress patterns. This is notably the case in hypocotyls, where the CMTs of the outer face of the epidermis are not aligned along the axis of maximum tension (i.e. perpendicularly to the axis of the hypocotyl) during the entirety of the growth phase (Chan *et al*, 2011; Crowell *et al*, 2011). Our comparative analyses of CMT organization and response to forces in the outer integument of the seed coat also show that the CMTs facing the different faces of outer integument cells do not necessarily organize according to predicted shape-driven stresses, even if they can still respond to the application of mechanical forces.

There may be several explanations for our results. The stresses experienced when seeds are compressed, or when an ablation is performed, could be different, both qualitatively and quantitatively, to those perceived when seed-coat walls are put under tension by the pressure of the expanding endosperm. This is consistent with recent work showing that CMTs respond differently to tensile forces and to compressive forces during hypocotyl-stretching experiments (Robinson & Kuhlemeier, 2018). It is also consistent with work in protoplasts showing that CMTs preferentially orient along the main stress direction when pressure is high, and to cell geometry when pressure is low (Colin *et al*, 2020). The stiffening of wall 3, which is an internal wall, could notably isolate the outer integument from the tension induced in the seed coat by endosperm expansion (Creff *et al*, 2015). This would explain why the CMTs of the outer side of the abaxial epidermis (facing wall 1) are organized during the anisotropic growth phase, when this wall is load-bearing, but not during the isotropic growth phase, when wall 3 becomes load-bearing. However, this does not explain why seed growth anisotropy apparently decreases before wall 3 starts to stiffen, or why the CMTs that face wall 2, in both abaxial and adaxial epidermis, remain organized at later stages of development, while the CMTs that face the load-bearing wall 3, are not particularly organized. Another possibility could be that tension is reduced in specific walls of the outer-integument as a result of changes in their thickness and/or stiffness caused by their differentiation. This possibility is supported by our immunolabelling studies, which show that antibodies targeting pectins with different degrees of methylesterification do not mark outer integument walls uniformly, and that the labelling also depends strongly on the developmental stage considered (Creff *et al*, 2023). It is also coherent with the literature showing that CMT organization and response to stresses can be enhanced by altering cell wall composition, for instance by inhibiting cellulose deposition in meristems (Heisler *et al*, 2010) or by altering pectin methylesterification in pavement cells (Tang *et al*, 2022). Nevertheless, our study supports the idea that CMT responses to stresses are key determinants of seed shape and that variations in these responses, in different outer integument layers, or at different stages of development, underlie the changes in growth anisotropy that are observed during seed development.

## Material and Methods

### Plant material and growth conditions

The *spr2-2* (CS6549 (Shoji *et al*, 2004)), *ktn1* (SAIL_343_D12 (Chen *et al*, 2014)), *ap2-6* (CS6241 (Wakem, 2003))*, iku2-2* (Garcia *et al*, 2005), *csi1-3* (SALK_138584 (Lei *et al*, 2014)), *prc1-1* (CS297 (Fagard *et al*)) and *ttl1 ttl3* (Salk_063943 and Sail_193_B05 (Lakhssassi *et al*, 2012)) mutants were previously described. The *p35S::Lti6b-GFP* (Cutler *et al*, 2000), *p35S::MAP65-1-RFP* (Creff *et al*, 2015), *p35S::TUA6-GFP* (Ueda et al, 1999), *pPDF1-MBD-mCitrine* (Malivert *et al*, 2021) and *pPDF1::CFP-N7* (Landrein *et al*, 2015) reporters were also described previously. The *pELA1::MAP65-mCitrine* reporter was developed for this study (see below). Seeds were sterilized in a solution of 70% ethanol and 0,05% Triton x100 (Sigma) for 15 min, rinsed 3 times in Ethanol 95%, and dried on Whatman paper for 30 minutes to 1h under the hood. Seeds were then sowed on plates containing 1X Murashige and Skoog (MS) medium (*pH* 5.7) and kept for two days in the dark at 4°C for stratification. They were then placed in a growth cabinet (Sanyo) under short-day condition (8h light, 21°C on daytime and 18°C on nighttime). After 2 weeks, germinated seedlings were transferred into individual pots of soil (Argile 10 (Favorit)), placed in a short-day room (8h light, 21°C on daytime and 19°C on nighttime) for a week before being transferred in a long day room for the rest of their lifecycle (16h light, 21°C on daytime and 19°C on nighttime). Note that the seeds used for the ablation experiment (Fig S6B) and the imaging of microtubules using the *pPDF1-MBD-mCitrine* line (Fig 5, S3 and S5) were grown differently. Following sowing on plates and stratification, they were placed in a long-day growth cabinet (16h light, 21°C on daytime and 19°C on nighttime) and transferred directly in a long day room (16h light, 21°C on daytime and 19°C on nighttime) after germination.

### Generation of the pELA1::MAP65-1-mCitrine line

The *pELA1* promoter was amplified and cloned into the pENTRY-R4-L1 as described in (Creff *et al*, 2015). A triple Gateway reaction (Life Technologies) was then performed using the *pELA1::pENTR-R4-L1* plasmid, a *MAP65-1-pENTR-L1-L2* plasmid and a *mCitrine-pENTR-R2-L3* plasmid as entry vectors and the *pS7m34GW* plasmid as destination vector to generate a *pELA1::MAP65-1-mCitrine-pS7m34GW* construct. Plants were transformed by flower-dipping using *Agrobacterium*. Transformants were selected based on their resistance to Sulfadiazine.

### Measurements of mature seed shape

Plants were grown under the conditions described in the previous section. Upon harvest, mature seeds were imaged using a SMZ18 Stereomicroscope (Nikon) mounted with a C11440 digital camera (Hamamatsu). The resulting black and white pictures were analysed using a dedicated macro on ImageJ. This macro uses a Huang thresholding to binarize the images and a distance transformed watershed algorithm to segment the seed contours and separate adjacent seeds. The seeds that were incorrectly segmented were manually removed from the analysis. For each seed, the aspect ratio was defined as the ratio between the major and the minor axis of the seed, which was automatically obtained using the “Fit Ellipse” option in ImageJ.

### Measurements of the size and shape of developing seeds

For all experiments but the reciprocal crosses and the one involving *ap2-6*, the seeds were staged every day for up to 10 days by marking the opening of the flowers on the main stem with coloured cotton threads. For the comparative analysis of Col-0 and *ap2-6* seed shape at 10 DPA and the reciprocal crosses, both WT and mutant flowers were emasculated before their opening and manually pollinated with Col-0 pollen to ensure that the seed growth defects were of maternal origin. The siliques were then harvested, put on double sided tape and opened with a needle so that their replum could be removed with forceps. The seeds on their replum were then put on a slide with a drop of clearing solution (1 vol glycerol/7 vol chloral hydrate liquid solution, VWR Chemicals). The seeds were detached from the replum and the slide was covered with a coverslip and kept in the dark for at least 24h in a cold room (4°C). The seeds were imaged with an Axioimager 2 (Zeiss) equipped with 10X and 20X DIC dry objectives and mounted with a Axiocam 705 color camera (Zeiss).

The measurements of seed size and shape were done using a semi-automatic Fiji macro. To measure seed area, the seed contour was extracted automatically using a Huang Dark auto-thresholding and a distance transformed watershed algorithm, or manually using the Polygon selection tool of ImageJ (for older seeds when the automatic segmentation was failing and for young seeds when the pedicel was still attached to the seed and could not be removed from the segmentation). The aspect ratio of the seed was defined as the ratio between the length and the width of the seed, which were measured manually using the straight line selection tool of ImageJ. The length of the seed was obtained by drawing a line from the pedicel of the seed to its tip that passes through the middle of the endosperm sac and its width was obtained by drawing a line perpendicular to the previous axis and passing through its centre.

### Live-imaging of developing seeds and analysis of cell growth rate and anisotropy

Plants expressing the ubiquitous membrane marker *p35S::LTi6b-GFP* were grown as described in previous sections and the opening of the flowers of the main stems was marked with a cotton thread. At either 1 or 4 days post-anthesis, the siliques were harvested and put on a slide covered with double-sided tape, and opened with a needle (under the hood, using sterilized material). The replum with its seeds attached was then extracted with tweezers and put on plates containing 2% Nitsch medium (Duchefa), 5% Sucrose (Sigma), 0.05% MES (Sigma), 1% agarose (Sigma), 0.5% Gamborg Vitamins (Sigma) and 0,1% Plant Preservative Mixture (PPM, Plant Cell Technology). Note that the medium (pH 5.8) needs to be autoclaved using a soft cycle (110°C for 10 minutes) and that the Gamborg vitamins and the PPM need to be added after autoclaving. Once put on the culture medium, the seeds, still attached to their replum, were covered with a drop of 0.5% Low Melting Agarose (Sigma) to prevent movement during imaging. The plates were then closed with plaster and placed in a long day growth cabinet (Panasonic, 16h light, 20°C) until imaging. To image the seeds, the plates were opened and covered with sterile water. Z-stacks (1µm thickness) were collected using a SP8 confocal microscope (Leica Microsystems) with a 25x long distance water-dipping objective (NA=0.95). The GFP was excited with a LED laser emitting at a wavelength of 488 nm (Leica Microsystems) and the signal between 495 nm and 555 nm was collected. After imaging of the first timepoint, the water was removed, the plates were closed, sealed again, and put back in the growth cabinet for 24h.

The resulting 3D stacks of developing seeds were analysed using the MorphographX software (Barbier de Reuille *et al*, 2015) following the protocol described in https://www.mpipz.mpg.de/4085950/MGXUserManual.pdf. The level set method (Kiss *et al*, 2017) was used to generate a 2.5D mesh of the surface of the seed on which the signal of the transverse walls of the abaxial epidermis was projected. To obtain the 2.5D surface of the adaxial epidermis, the surface obtained for the abaxial epidermis by the level set method was translated in z of -10 µm and the level set method (option “evolve”) was used again for several cycles to detect the longitudinal wall separating the abaxial from the adaxial epidermis (wall 2). A 2.5D mesh of this new surface was then generated, on which the signal of the transverse walls of the adaxial epidermis was projected. Note that this method generates a 2.5D mesh of the surface of the adaxial epidermis that is of a lower quality than the one of the abaxial epidermis. In both layers, the cells were then manually seeded and their contour was segmented using the watershed algorithm of the software. The parent to offspring association (lineage) of the cells of two successive timepoints was done manually according to the software guide. Heatmaps of cell size, aspect ratio, growth rate and growth anisotropy were then generated according to the software guide. The data were exported as csv files and analysed on R (https://www.r-project.org/). The graphs displaying growth rate and anisotropy as a function of the position of the cell along the main axis of the seed were generated as described in: (Strauss *et al*, 2022).

### Measurements of microtubule organization in the seed outer integument layers

The *p35S::MAP65-1-RFP*, *pPDF1-MBD-mCitrine* and *p35S::TUA6-GFP* reporters were used to image the CMTs of the outer face of the abaxial outer integument epidermis. The *pPDF1-MBD-mCitrine* was also used to compare CMT organization in both faces of the abaxial epidermis, being only expressed in this layer. The *pELA1::MAP65-1-mCitrine* reporter was used to compare CMT organization in both faces of the adaxial epidermis, being only expressed in this layer.

The siliques were opened and the seeds were placed individually on adhesive tape on a microscope slide and covered with water. All acquisitions were performed using a Zeiss 980 Airyscan microscope with a water-dipping plan apochromat 20x objective (NA=1), except for the seed compressions at 2 DPA (Fig 4) where a SP8 confocal microscope (Leica Microsystems) with a 25x long distance water-dipping objective (NA=0.95) was used, and the cell ablations and the imaging of the CMTs facing of the abaxial epidermis using the *pPDF1::MBD-mCitrine* line (Fig S3 and S6), where a Nikon C2 with a 60x water-dipping objective was used (NA=1). The GFP was excited with a LED laser emitting at a wavelength of 488 nm (Leica Microsystems) and the signal between 495 nm and 555 nm was collected. The RFP was excited with a LED laser emitting at a wavelength of 552 nm and the signal was collected at 580-650 nm. The mCitrine was excited with a LED laser emitting at a wavelength of 514 nm and the signal was collected at 520-560 nm.

The mean orientation and organization of the CMTs per cell were all measured in 2D in the flat part of the growing area of the seed using the FibrilTool macro in ImageJ (Boudaoud *et al*, 2014), except for the measurements of CMT organization following seed compression at 2DPA (Fig 4) and for the measurements of the correlation between CMT orientation and local seed curvature, which were done on 2.5D meshes using the FibriTool macro integrated in the MorphographX software. Note that the degree of CMT organization (also referred as the degree of anisotropy) is obtained using a nematic tensor and is dependent on the nature of the CMT reporter that is used but also on the quality of the picture. This explains why the degree of organization of the CMTs obtained using ImageJ is lower than the one obtained using MorphographX as the first one only considers the signal of few slices while the second uses the signal of many slices projected on the 2.5D surface of the seed. Also note that the brightness and contrast of the CMTs images were enhanced for a better visualization of their organization.

To generate the heatmaps and frequency plots showing the correlation between the orientation of the CMTs in the outer face of the abaxial epidermis and the local direction of maximum curvature at each cell position, maps of the mean Gaussian curvature, projected on the cell-segmented 2.5D meshes, were generated with MorphographX. Given that the cells in the abaxial epidermis enlarge from 2 to 5 days post-anthesis (DPA), a neighbouring of 30 µm was used to generate the curvature maps at 2 DPA while a neighbouring of 60 µm was used to generate the curvature maps at 5 DPA. This neighbouring roughly corresponds, in both cases, to the distance between the centre of the considered cell and the furthest edge of its direct neighbours. The angle θ representing the difference between CMT orientation (defined as a vector (xMt, yMt, zMt)) and maximum curvature direction (defined as a vector (xMc, yMc, zMc)) was calculated using the following formula:

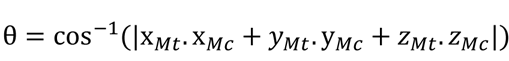

Note that both vectors are tangent to the surface of the mesh at the centre of each cell considered.

### Application of mechanical constraints to developing seeds

For the compression, the entire silique was compressed using a microvice as described in (Creff *et al*, 2015) and kept for 24h in a growth chamber before preparation of the seeds and subsequent imaging of the CMTs as described in the previous sections. Optical sections of the 3D stacks were generated to check that the seeds were efficiently compressed. For the ablation, the seeds were prepared as described in the live-imaging section. The ablation was done manually using a sharp needle under a stereomicroscope as described in: (Uyttewaal *et al*, 2012). Seeds were kept in a closed box between two timepoints. The measurements of the orientation of the CMTs in each cell relative to the position of the ablation was done in 2D on ImageJ as described in (Malivert *et al*, 2021).

### Statistical analysis and reproducibility

All experiments except the measurements of the shape of mature seeds in the different CMT reporters, have been carried at least two times independently and the resulting data have been pooled. The plots were generated using the R software, Python or Excel. When data are compared using Student tests, statistical significance was displayed with stars as followed: *p<0.05, **p < 0.01, ***p<0.001 and ****p < 0.0001. In the boxplot representations, the midline represents the median of the data while the lower and upper limits of the box represent the first and third quartile respectively. The error bars represent the distance between the median and one and a half time the interquartile range. When possible, individual values, corresponding to single seeds, are also overlayed as points. When relevant, violin plots are also displayed to get a better grasp of the shape of the distribution. In the frequency plots, the bold lines show the trending curves and the dotted lines show the mean value when all cells and seeds of a given condition are pooled. The plots showing seed growth rate and seed growth anisotropy were obtained by deriving the daily measurements of seed size and aspect ratio using the following formula:

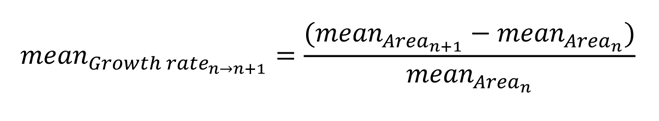

The error bar show the standard deviation calculated using the following formula:

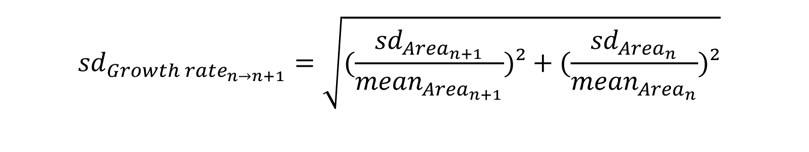

## Acknowledgements

We thank Stéphane Verger, Alice Malivert and Olivier Hamant for providing the *ktn1*, *csi1-3, prc1-1* and *spr2-2* mutants and the *p35S::TUA6-GFP, pPDF1::mCitrine-MBD and pPDF1::CFP-N7* reporters, Miguel Botella for providing the *ttl1 ttl3* mutant, Frederic Berger for providing the *iku2-2* mutant, Marilyn Vantard for providing the *p35S::MAP65-1-RFP* reporter and Marie-Cecile Caillaud for providing the *MAP65-1-pENTR-L1-L2* plasmid. We also thank Olivier Hamant, Marie-Cécile Caillaud, Magalie Uyttewaal, Olivier Ali, and Charlotte Kirchhelle for helpful discussion and comments on the manuscript; Audrey Creff, Vincent Bayle, Olivier Ali, Claire Lionnet and Corentin Mollier for technical assistance with experiments and analysis; the PLATIM (*SFR128 Biosciences* Lyon) and the Bio21 Advanced Microscopy Facility (University of Melbourne) for technical assistance with microscopy; Alexis Lacroix, Patrice Bolland and Justin Berger for technical assistance with plant cultivation; Isabelle Desbouchages and Hervé Leyral for technical assistance regarding molecular biology work; Cindy Vial, Laureen Grangier, Nelly Camilleri and Stéphanie Maurin for administrative assistance. Amélie Bauer was supported by a joint PhD program between the CNRS and the University of Melbourne. Camille Bied is supported by a PhD fellowship from the French Ministry of Higher Education.

## Author contributions

B.L., J.G. and G.I. led the study, obtained funding and supervised the work. A.B., C.B., A.D. and B.L. carried and analysed the experiments. A.B. and B.L. wrote the paper with input from all co-authors.

## Data and code availability

All the raw data relative to this study will be deposited on the Zenodo repository upon publication. The code specifically developed for this study will be deposited on Github.

## Supplementary Figures

**Fig. S1.**
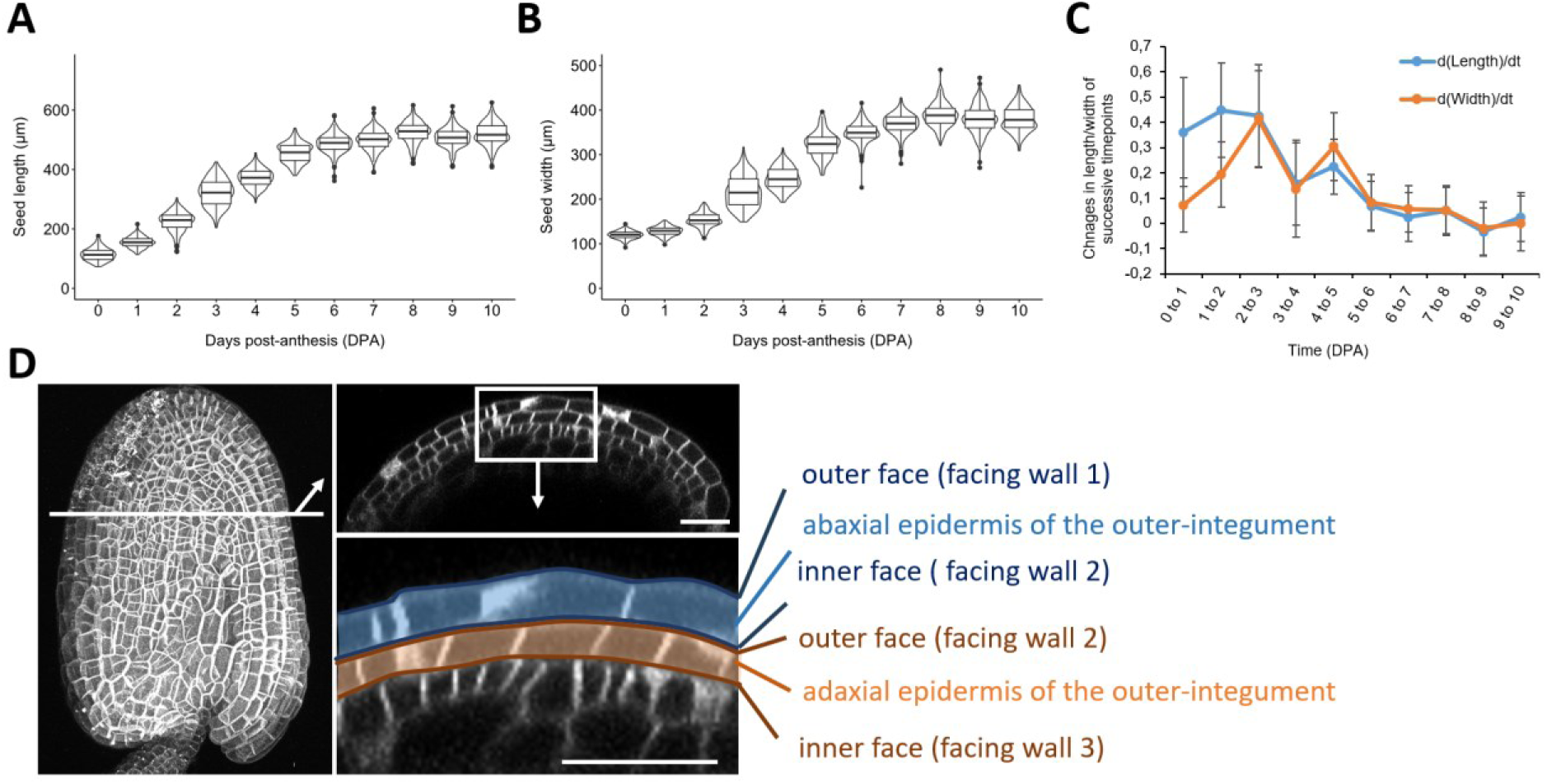
Characterization of the structure and growth pattern of the seed. **A.** Measurements of the length of WT seeds (Col-0 ecotype) from 0 to 10 days post-anthesis (10 DPA), n=180-209 seeds per day, two independent experiments. **B.** Measurements of the width of WT seeds (Col-0 ecotype) from 0 to 10 days post-anthesis (10 DPA), n=180-209 seeds per day, two independent experiments. **C.** Relative changes in seed length and width over time obtained by deriving the measurements of seed length and width of panels A and B. **D.** Z-projection, and optical section passing through the middle of a seed at 2 DPA expressing the ubiquitous membrane marker *LTi6b-GFP* showing the organization of the seed coat. The two outermost cell layers of the seed coat are the abaxial and the adaxial epidermis of the outer integument. In each of these layers, we defined the longitudinal wall facing the outer side of the seed as the “outer face” and the longitudinal wall facing the inner side of the seed as the “inner face”. These walls can also be numbered based on their position in regards to the outside of the seed (from 1 to 3), wall 1 being the outermost-cell wall of the seed coat and wall 3 being the wall at the interface between the inner and the outer integument. Scale bars: 20 µm.

**Fig. S2.**
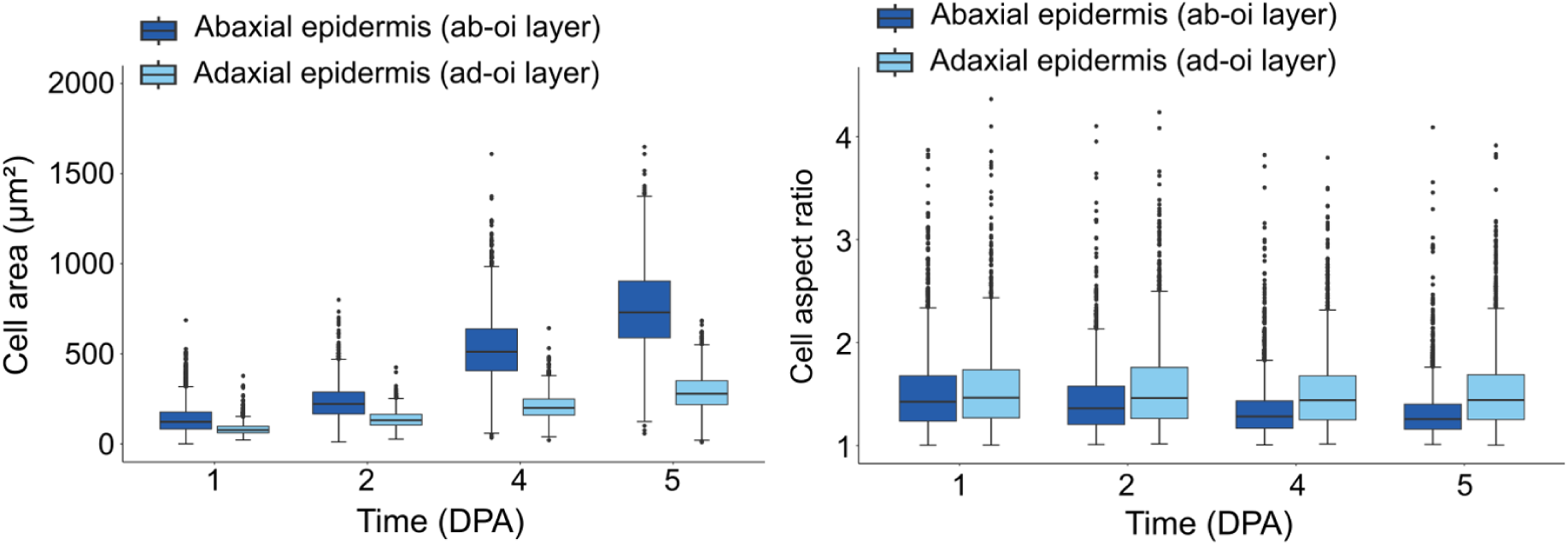
Cells in the outer integument change more in size than in shape during seed development. Evolution of the size (measured as area) and shape (measured using the aspect ratio) of the cells in the abaxial and adaxial epidermis of the outer integument (DPA: Days post-anthesis), 1373 to 2402 cells from 10 to 11 seeds, two independent experiments.

**Fig. S3.**
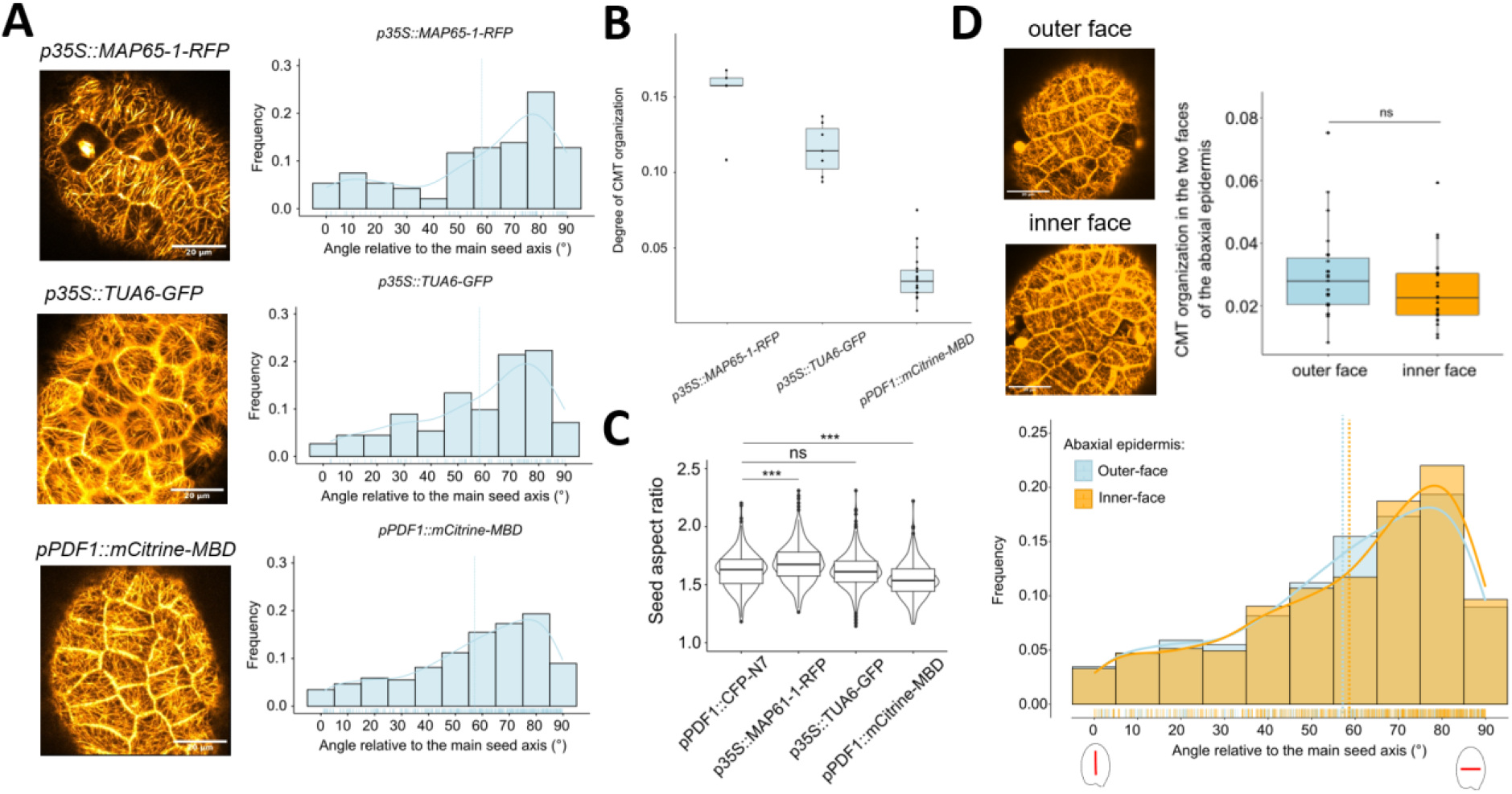
Different CMT reporters show similar orientations of the CMTs in the abaxial epidermis but different levels of organization, which correlate with changes in seed shape. **A.** Orientation of the CMTs facing the outer side of the seed in the abaxial outer integument epidermis at 2 days post anthesis (DPA) and assessed using three different CMT reporters: *p35S::MAP65-1-RFP, p35S::TUA6-GFP* and *pPDF1::mCitrine-MBD*, scale bars: 20 µm, n= 94 to 491 cells from 5 to 23 seeds, two to three independent experiments depending on the reporter. In the frequency plots, the bold lines show the trending curves and the dotted lines show mean orientations relative to the seed axis. **B.** Mean degree of organization of the CMTs in the outer-face of the abaxial epidermis per seed at 2 DPA obtained using three different CMT reporters: *p35S::MAP65-1-RFP, p35S::TUA6-GFP* and *pPDF1::mCitrine-MBD*, n= 94 to 491 cells from 5 to 23 seeds, two to three independent experiments depending on the reporter. **C.** Aspect ratio of mature seeds of a control line (*pPDF1::CFP-N7*) and of the three CMT reporters presented in A and B, n=349-554 seeds, one experiment. Data were compared using Bilateral Student tests. The aspect ratio of mature seeds positively correlates with the degree of organization of the CMTs in the outer face of abaxial outer integument epidermis measured during the anisotropic growth phase. **D.** Comparison between the mean degree of organization (top) and the mean orientation (bottom) of the CMTs (*pPDF1::mCitrine-MBD* reporter) facing the outer side and inner side of the seed in the abaxial epidermis at 2 DPA, scale bars: 20µm, n= 491 cells from 23 seeds, three independent experiments. Data were compared using Bilateral Student tests. In the frequency plot, the bold lines show the trending curves and the dotted lines show mean orientations relative to the seed axis.

**Fig. S4.**
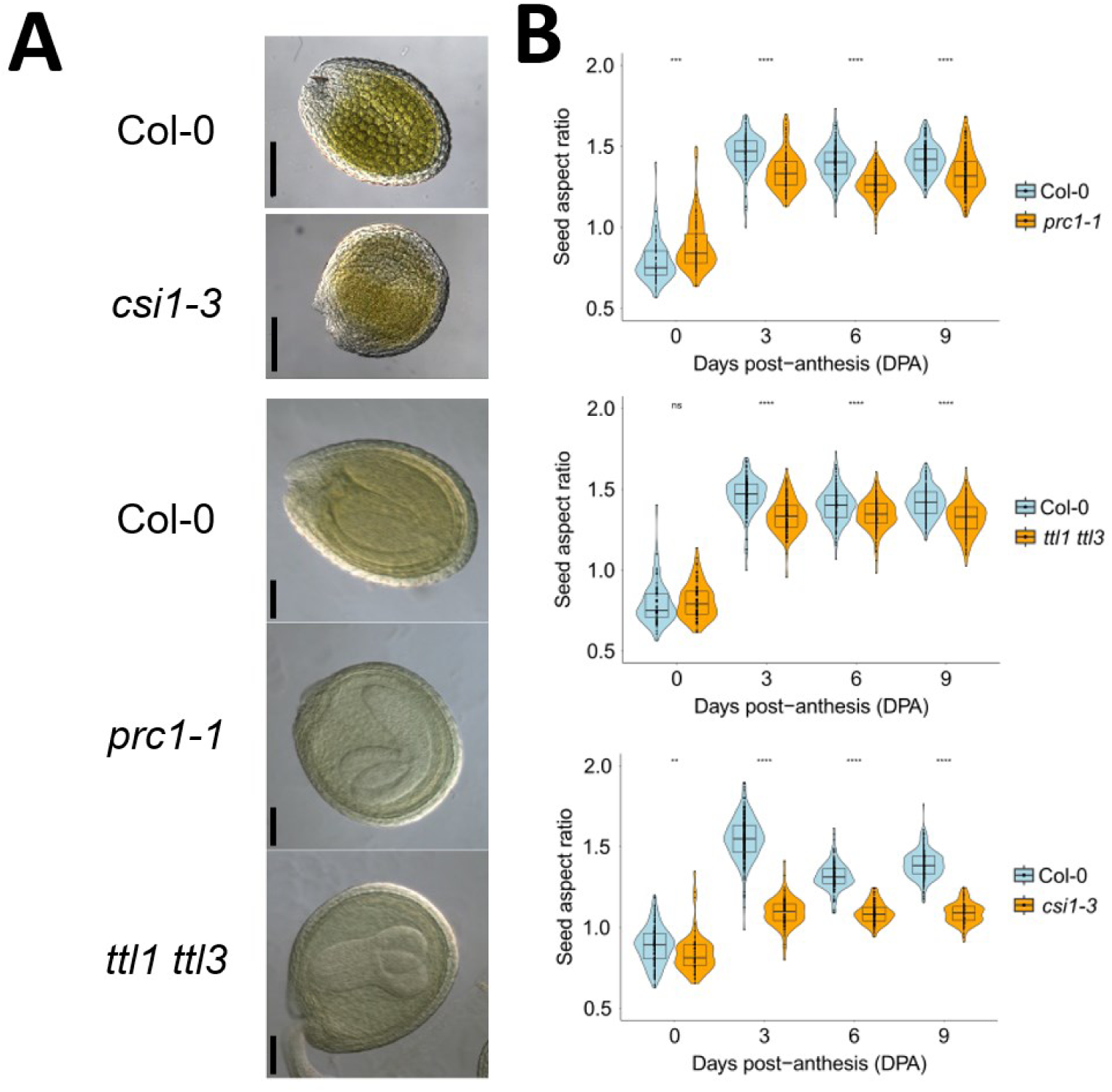
Altering CMT-guided cellulose deposition affect seed growth anisotropy. **A.** Seed shape defects of the *csi1-3, prc1-1 (cesa6)* and *ttl1 ttl3* mutants at 10 days post-anthesis (DPA). Scale bars: 0.1 mm. **B.** Aspect ratio of developing WT and *csi1-3, prc1-1 (cesa6)* and *ttl1 ttl3* mutant seeds, n= 69 to 366 seeds per day, two independent experiments. Data were compared using bilateral Student tests.

**Fig. S5.**
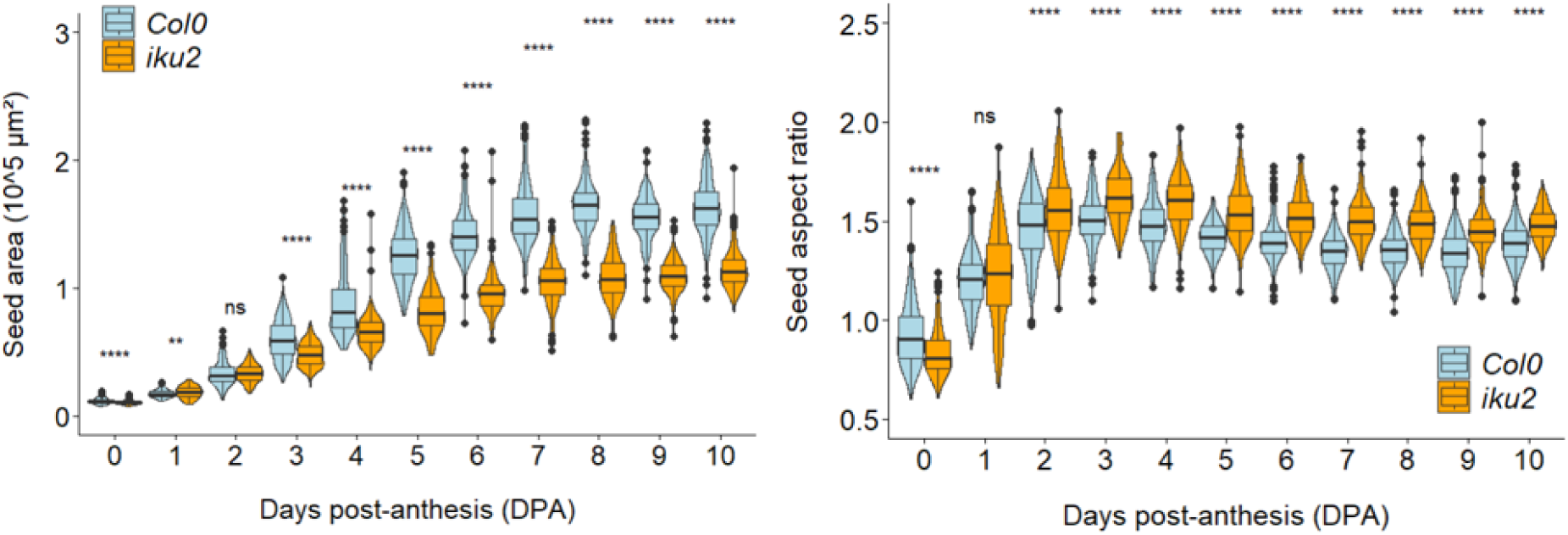
Seed size and shape are altered in the endosperm-pressure mutant *iku2*. Measurements of the area and aspect ratio of developing seeds from 0 to 10 days post-anthesis (DPA) in the Wild-Type (Col-0) and the endosperm-pressure defective mutant *iku2*., n= 207-313 seeds per day per genotype, three independent experiments. Data were compared using bilateral Student tests.

**Fig. S6.**
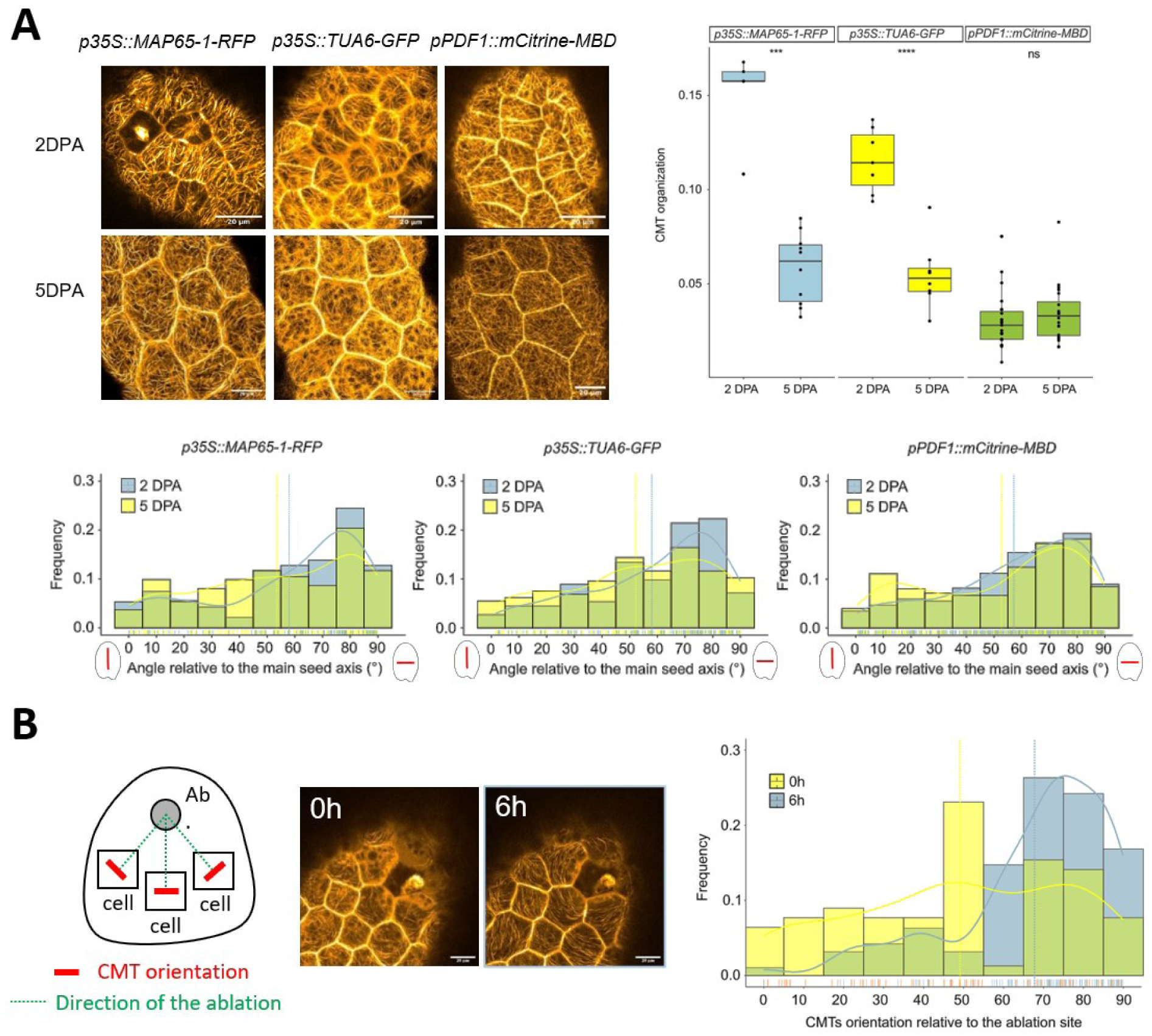
The CMTs facing the outer face of the abaxial outer integument epidermis lose their organization but not their mechanosensitivity during the transition from anisotropic to isotropic growth. **A.** Organization and orientation of the CMTs in the outer face of the abaxial outer integument epidermis during the anisotropic growth phase (2DPA) and during the isotropic growth phase (5DPA) assessed using three different CMT reporters (*p35S::MAP65-RFP, p35S::TUA6-GFP*, *pPDF1::mCitrine-MBD*), scale bars: 20µm, n= 94 to 491 cells from 5 to 23 seeds, two to three independent experiments depending on the reporter. Data were compared using bilateral Student tests. In the frequency plots, the bold lines show the trending curves and the dotted lines show mean orientations relative to the seed axis. **B.** Effect of a cell ablation on the orientation of the CMTs (imaged using the *p35S::MAP65-1-RFP* reporter) facing the outer side of the seed in the abaxial outer integument epidermis, n= 78 to 95 cells of 12 to 13 seeds, three independent experiments. In the frequency plot, the bold lines show the trending curves and the dotted lines show mean orientations of the CMTs relative to the position of the ablation.

